# Aliphatic Amines are Viable Pro-drug Moieties in Phosphonoamidate Drugs

**DOI:** 10.1101/2020.04.05.026583

**Authors:** Victoria C. Yan, Cong-Dat Pham, Kenisha Arthur, Florian L. Muller

**Affiliations:** Department of Cancer Systems Imaging, University of Texas MD Anderson Cancer Center, Houston, TX, 77054, USA

**Author notes:** Correspondence should be addressed to Florian L. Muller, Department of Cancer Systems Imaging, University of Texas MD Anderson Cancer Center, Houston, TX 77054.

## Abstract

Phosphate and phosphonates containing a single P-N bond are frequently used pro-drug motifs to improve cell permeability of these otherwise anionic moieties. Upon entry into the cell, the P-N bond is cleaved by phosphoramidases to release the active agent. Here, we apply a novel mono-amidation strategy to our laboratory’s phosphonate-containing glycolysis inhibitor and show that a diverse panel of phosphonoamidates may be rapidly generated for in vitro screening. We show that, in contrast to the canonical L-alanine or benzylamine moieties which have previously been reported as efficacious pro-drug moieties, small aliphatic amines demonstrate greater drug release efficacy for our phosphonate inhibitor. These results expand the scope of possible amine pro-drugs that can be used as second pro-drug leave groups for phosphate or phosphonate-containing drugs.

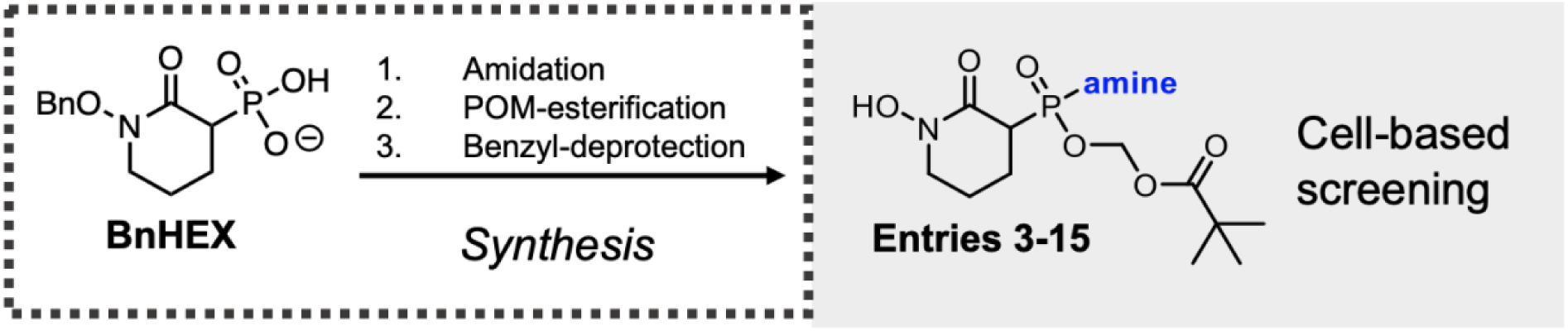

---

Phosphoramidates are structurally intriguing chemical moieties with high biological and therapeutic relevance^1^. Within the realm of drug development, the inclusion of a phosphoramidate moiety has become an increasingly attractive pro-drug strategy for anionic phosphate^2,3^- or phosphonate^4–6^-containing drugs. Upon entering the cell, the P-N bond would eventually be cleaved by a class of enzymes known as phosphoramidases^7^; this includes a family of enzymes known as histidine triad nucleotide-binding proteins (HINT1, HINT2, HINT3)^8^. From anti-viral drugs such as Tenofovir alafenamide^9^, Sofosbuvir^10^, and Remdesivir^11^ to emerging pro-drugs of conventional chemotherapies such as Gemcitabine (Acelarin^12^) and 5-Fluorouracil (NUC-3373, NCT03428958^13^), the phosphoramidate-containing “ProTide”^14^ approach to delivering phosphate or phosphonate drugs is a central theme in pro-drug development^1,15^. Common to these drugs is the presence of an L-alaninate ester moiety, which is used as a substrate for phosphoramidase cleavage. To the best of our knowledge, apart from the L-alaninate ester, the only other amine that has been reported in the context of phosphoramidate or phosphonoamidate pro-drugs is the benzylamine moiety found in IDX-184^16^. These observations would suggest that either the L-alaninate or benzylamine moieties would be the most optimal amine substrates for release by phosphoramidases. Fundamental biochemical studies have explored the relationship between altering the amine on phosphoramidate versions of AMP rate of hydrolysis by the phosphoramidases in vitro^7,8,17^. These studies suggest that amines beyond L-alaninate esters and benzylamine may also be used as second pro-drug groups. Perhaps due to the difficulty in preparing mono-amidated substrates, however, further investigation into the structural limits of optimal amine leave groups in other contexts beyond adenine monophosphate have yet to be reported. Here, we report the efficacy of various structurally diverse amine pro-drug groups in the context of our laboratory’s phosphonate inhibitor of the glycolytic enzyme Enolase, termed HEX^18^. We validate the finding that benzylamine and other benzylic amines may be used as second pro-drug groups and also demonstrate that, in the context of HEX, aliphatic amines are often superior delivery moieties. These results may prompt further investigation into using aliphatic amines as pro-drug moieties for phosphate or phosphonate-containing drugs.

We generated a diverse panel of phosphonoamidate pro-drugs of HEX using a novel mono-amidation reaction that we had previously discovered (Yan 2020, manuscript under review). Identifying amines with superior pro-drug properties is a central focus in our laboratory’s endeavor towards optimizing the delivery of our active Enolase inhibitor, HEX. We previously described an innovative therapeutic strategy, called collateral lethality, wherein cancer cells harboring the homozygous deletion of an essential metabolic gene are selectively sensitized to inhibition of its redundant paralogue^5,19^. Inhibition of the ENO2 isoform of Enolase in *ENO1*-deleted cancers results cancer-specific cell death—leaving *ENO1*-intact or normal cells unperturbed^18^. To evaluate the relationship between our amine pro-drug structure and efficacy of drug delivery, we first esterified the other free hydroxyl on the phosphonic acid to enable efficient cell permeability (**Figure 1a**). We chose the pivaloyloxymethyl (POM) group as the first ester group due to its known susceptibility for hydrolysis by ubiquitously expressed carboxylesterases^18,20^. Importantly, our decision to employ the labile POM ester minimizes the confounding effects of initial pro-drug cleavage and ensures that relationship between amine structure and cell killing can be studied in isolation (**Figure 3b**). The mono-amidated products generated from our reactions with BnHEX were subject to POM esterification, followed by hydrogenation to liberate the hydroxamate—an absolutely essential moiety for Enolase active site inhibition^5,18^.

**Figure 1.**
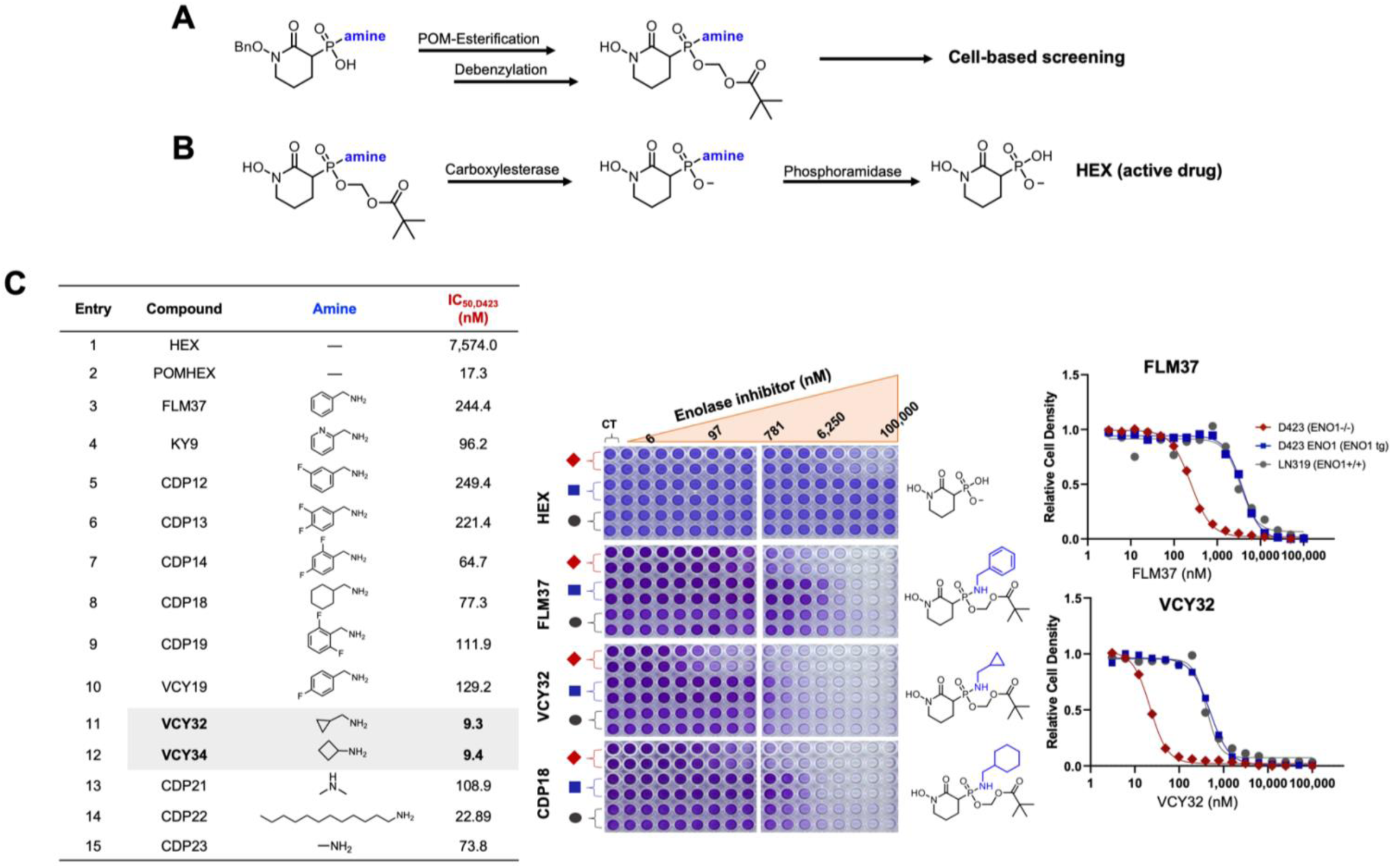
Aliphatic amines offer superior drug delivery in vitro. (**a**) General workflow for the synthesis and screening amidated pro-drugs. For BnHEX, POM esterification followed by de-benzylation of the hydroxamate yields the final, cell-permeable pro-drug. For specific reaction conditions, see Supplementary Information. (**b**). Proposed bioactivation mechanism for phosphonoamidate pro-drugs. Phosphoramidases cleave P-N bonds on anionic molecules^7^ and can thus serve as second pro-drug deprotecting enzymes. (**c**) The relationship between amine structure and pro-drug efficacy can be evaluated in cell-based screening. (Left) Structures of amine pro-drugs of the Enolase inhibitor, HEX, and corresponding IC_50_s against D423 (*ENO1*-deleted) cells. (Middle) Crystal violet cell proliferation assay evidences greater cell killing against D423 cells by aliphatic amine pro-drugs compared to benzylamine. Cells were incubated with pro-drug inhibitor for 5 days. Then, cells were fixed and stained with crystal violet and quantified spectroscopically. Cell density as measured by crystal violet were plotted as a function of inhibitor. (Right) Comparison of the IC_50_ values between model aromatic (FLM37) and aliphatic (VCY32) pro-drugs. While both pro-drugs are selective for *ENO1-*deleted cells, VCY32 exhibits 10-fold greater potency compared to FLM37 (IC_50_ = 22 nM versus 244 nM).

Esterified phosphonoamidates all demonstrated selective, nanomolar activity against *ENO1*-deleted glioma cells (D423) compared to *ENO1*-rescued (D423 ENO1) and *ENO1*-wildtype glioma cells (LN319) (**Figure 1c**). Given the focus on aryl and benzylic amines as pro-drug motifs in the literature^7,21^, we were surprised to find that aliphatic amines proved to be most effective. Cyclopropylmethanamine-protected HEX exhibited 10-fold greater potency compared to benzylamine-protected HEX (IC_50, D423_ = 22 nM versus 244 nM, **Figure 1c**). Direct comparison of benzylamine-protected HEX to its saturated counterpart, cyclohexane methylamine, likewise showed greater potency of the latter (**Figure 1c, entry 3 versus 8**). Apart from these more striking examples, our data suggest that mono-substitution at the *ortho* position of the benzylamine ring can enhance cell killing ability and, by extension, hydrolytic activity by phosphoramidases (**Figure 1c, entries 4**,**7**,**9**). Substitution at the *meta* position does not seem to improve the properties of benzylic ring, as observed for the nearly identical IC_50_s between benzylamine and 3-fluorobenzylamine (**Figure 1c, entry 2 versus 5**). Collectively, these data show that fluorine substitution on the aromatic ring is well-tolerated, pointing to the feasibility of applying ^18^F-labeling methods to study pharmacodynamics for any phosphoramidase-labile pro-drug. More generally, we have demonstrated that aliphatic amines—especially those with low molecular weights—are viable pro-drug moieties and can offer superior delivery of phosphonate compounds in vitro **(Figure 1, entries 11-15**). Taken together, these results point to the viability of using (small) aliphatic amines as pro-drug moieties for phosphates or phosphonates. Compared to L-alaninate esters or benzylamine, small aliphatic amines have lower molecular weights, which could facilitate more efficient cell permeability^22^. Thus, our findings expand the scope of possible amine pro-drugs that can be used as second pro-drug leave groups.

## AUTHOR CONTRIBUTIONS

C.D.P. and V.C.Y performed syntheses and characterized compounds. K.A. performed cell culture experiments. V.C.Y. wrote the manuscript. F.L.M. provided critical comments and oversaw the project.

## ACKNOWLEDGEMENTS

This work is supported by the American Cancer Society (RSG-15-145-01-CDD) and the National Comprehensive Cancer Network (YIA170032).

